# CD39 expression by regulatory T cells drives CD8+ T cell suppression during experimental *Trypanosoma cruzi* infection

**DOI:** 10.1101/2023.09.14.557792

**Authors:** Cintia L. Araujo Furlan, Santiago Boccardo, Constanza Rodriguez, Simon C. Robson, Adriana Gruppi, Carolina L. Montes, Eva V. Acosta Rodríguez

## Abstract

An imbalance between suppressor and effector immune responses may preclude cure in chronic parasitic diseases. In the case of *Trypanosoma cruzi* infection, specialized regulatory Foxp3+ T (Treg) cells suppress protective type-1 effector responses. Herein, we investigated the kinetics and underlying mechanisms behind the regulation of protective parasite-specific CD8+ T cell immunity during acute *T. cruzi* infection. Using the DEREG mouse model, we found that Treg cells play a critical role during the initial stages after *T. cruzi* infection, subsequently influencing CD8+ T cells. Early Treg cell depletion increased the frequencies of polyfunctional short-lived, effector T cell subsets, without affecting memory precursor cell formation or the expression of activation markers. In addition, Treg cell depletion during early infection minimally affected the antigen-presenting cell response but it boosted CD4+ T cell responses before the development of anti-parasite effector CD8+ T cell responses. Crucially, the absence of CD39 expression on Treg cells significantly bolstered effector parasite-specific CD8+ T cell responses, leading to improved parasite control during *T. cruzi* infection. Our work underscores the crucial role of Treg cells in regulating protective anti-parasite immunity and provides evidence that CD39 expression by Treg cells represents a key immunomodulatory mechanism in this infection model.

## Introduction

The protozoan parasite *Trypanosoma cruzi* is the causative agent of Chagas disease, a neglected tropical disease, which is endemic to South and Central America, with some regions of North America also impacted. Due to human migration, Chagas disease has become a worldwide concern (1). Early detection of infection is challenging and access to curative, non-toxic pharmacological treatment is limited, often resulting in lifelong disease for most individuals. During the acute phase, *T. cruzi* undergoes intensive replication, becoming detectable in the bloodstream and then spreading systemically to host tissues. The infection triggers a strong type 1 pro-inflammatory response, involving activation of cells from both the innate and adaptive immune system (2).

Macrophages, neutrophils, natural killer cells and dendritic cells act as the first defense against *T. cruzi,* while specific T and B cell responses develop over time. Notably, highly immunodominant parasite-specific CD8+ T cell responses play a crucial albeit delayed role in the optimal control of parasite replication (3, 4). Immune help from CD4+ T cells is required to achieve robust parasite-specific CD8+ T cell responses (5). However, although effector mechanisms are essential for limiting systemic acute infection, the immune response often fails to completely eliminate *T. cruzi*, allowing pathogen persistence in target tissues and leading to chronic disease. While a substantial proportion of patients remain asymptomatic during the chronic phase, approximately 30-40 % of infected individuals develop life-threatening cardiac and/or gastrointestinal complications.

Regulatory T (Treg) cells are a distinct subset of CD4+ T cells that are characterized by the expression of the transcription factor Foxp3, which equips them with the remarkable capacity to exert suppressive effects on a wide range of immune cell populations (6). During infectious inflammatory responses, Treg cell-mediated immunity may develop once the infection has been controlled to restore homeostasis and/or prevent collateral tissue damage. Conversely, pathogens can also induce Treg cells, as an early mechanism of immune evasion. As a result, Treg cells can have both beneficial and detrimental effects during infections, demanding a delicate and kinetic balance between effector and regulatory responses to achieve pathogen clearance, while protecting against excessive inflammation (7). Treg cell accumulation has been observed in peripheral blood, secondary lymphoid organs and target tissues of chronic bacterial, viral and parasitic infections in both mice and humans (6, 8). Moreover, decreased numbers of Treg cells have been reported during the acute phase of a few infections, such as those caused by *Toxoplasma gondii* (9, 10), *Listeria monocitogenes* (9), vaccinia virus (9), LMCV clone Armstrong (11), and more recently, Chikungunya virus (12). These findings underscore the complexity of Treg cell dynamics and illustrate the context-dependent roles in different infectious scenarios.

We have previously described a relative decrease in Treg cell numbers during acute *T. cruzi* infection, which, unlike most infections, did not fully recover in the transition to the chronic phase (13). Furthermore, we discovered that once Treg cells became activated, these cells acquired both phenotypic and transcriptional profiles consistent with suppression of type 1 inflammatory responses. Our findings have also highlighted the critical role of the Treg cell response natural contraction in allowing the emergence of protective anti-parasite CD8+ T cell immunity, as noted during the acute phase of *T. cruzi* infection. In line with these observations, a study by Ersching *et al.* reported that Treg cells suppress CD8+ T cell priming in an immunization model using *T. cruzi*-stimulated dendritic cells (14). Through the use of immune deficient mice and blocking antibody treatments, these authors have demonstrated that this effect was independent of IL-10 but partially mediated by CTLA-4 and TGF-β. However, the specific mechanisms employed by Treg cells to suppress effector responses during acute *T. cruzi* infection have yet to be fully elucidated.

CD39, also known as ectonucleoside triphosphate diphosphohydrolase 1 (ENTPDase1/gene *ENTPD1*), is a nucleotide-metabolizing enzyme involved in immune regulation and inflammation (15). In response to pro-inflammatory stimuli, intracellular ATP is released into the extracellular space, acting as a danger signal that promotes inflammation. However, the presence of CD39-expressing “regulatory” cells modulate this process by catalyzing ATP to AMP conversion. In turn, AMP can be further converted into the immunoregulatory molecule adenosine by CD73. Thus, the tandem activity of CD39 and CD73 holds the potential to shift the inflammatory environment towards an immunosuppressive state through the degradation of ATP into adenosine (16). CD39 expression has been identified in various immune cell populations, including Treg cells, in both mice and humans – as well as marking immune exhaustion (17–19). Consequently, the investigation of Treg cells expressing CD39 in different inflammatory contexts has shed light on their significant contributions to autoimmune diseases, cancer, allergies, and viral infections (20, 21). However, the role of CD39+ Treg cells remains largely unexplored in the context of microbial and parasitic infections, warranting further investigation to elucidate their potential implications in such settings.

Herein, we aimed to comprehensively investigate the biological significance of Treg cells in the context of *T. cruzi* infection. We sought to determine the specific time window during which Treg cells exert immune suppression and to identify the immune cell populations that are particularly sensitive to these regulatory effects in the acute phase of this parasitic infection. Through our investigations, we uncovered that during the initial stages of infection, endogenous Treg cells play a critical and unwelcome role in suppressing the expansion of CD4+ T cells and the development of anti-parasite effector CD8+ T cell responses. Moreover, we discovered that CD39 serves as a crucial regulatory mechanism through which Treg cells mediate suppression of *T. cruzi*-specific CD8+ T cells. These findings provide valuable insights into the role of Treg cells during the acute phase of *T. cruzi* infection and suggest potential avenues for immunomodulatory therapeutic strategies targeting CD39 expression and function.

## Results

### Treg cell depletion expands parasite-specific CD8+ T cells and improves parasite control during acute *T. cruzi* infection

We previously demonstrated that an exogenous increase in Treg cell numbers during acute *T. cruzi* infection dampen the anti-parasite effector response and subsequently influences infection control (13). To further investigate the biological role of the endogenous Treg cell response and elucidate the underlying mechanisms involved in effector T cell suppression during this infection, we employed the DEREG (DEpletion of REGulatory T cells) mouse model (22) to conduct Treg cell depletion experiments.

As illustrated in Figure 1A, DEREG mice were infected and received diphtheria toxin (DT) injections at days 5 and 6 post-infection (pi), in order to target Treg cells before the emergence of adaptive anti-parasite responses that occurs at day 10 pi in our infection model (23). We observed that the depletion strategy efficiently reduced Treg cell frequencies in the blood of DT-treated mice compared to PBS-treated controls, starting from at least day 11 pi (Figure 1B). A kinetics analysis in blood revealed that Treg cell frequencies returned to normal levels around 15 days after DT treatment in non-infected (NI) mice (Figure 1C).

**Figure 1.**
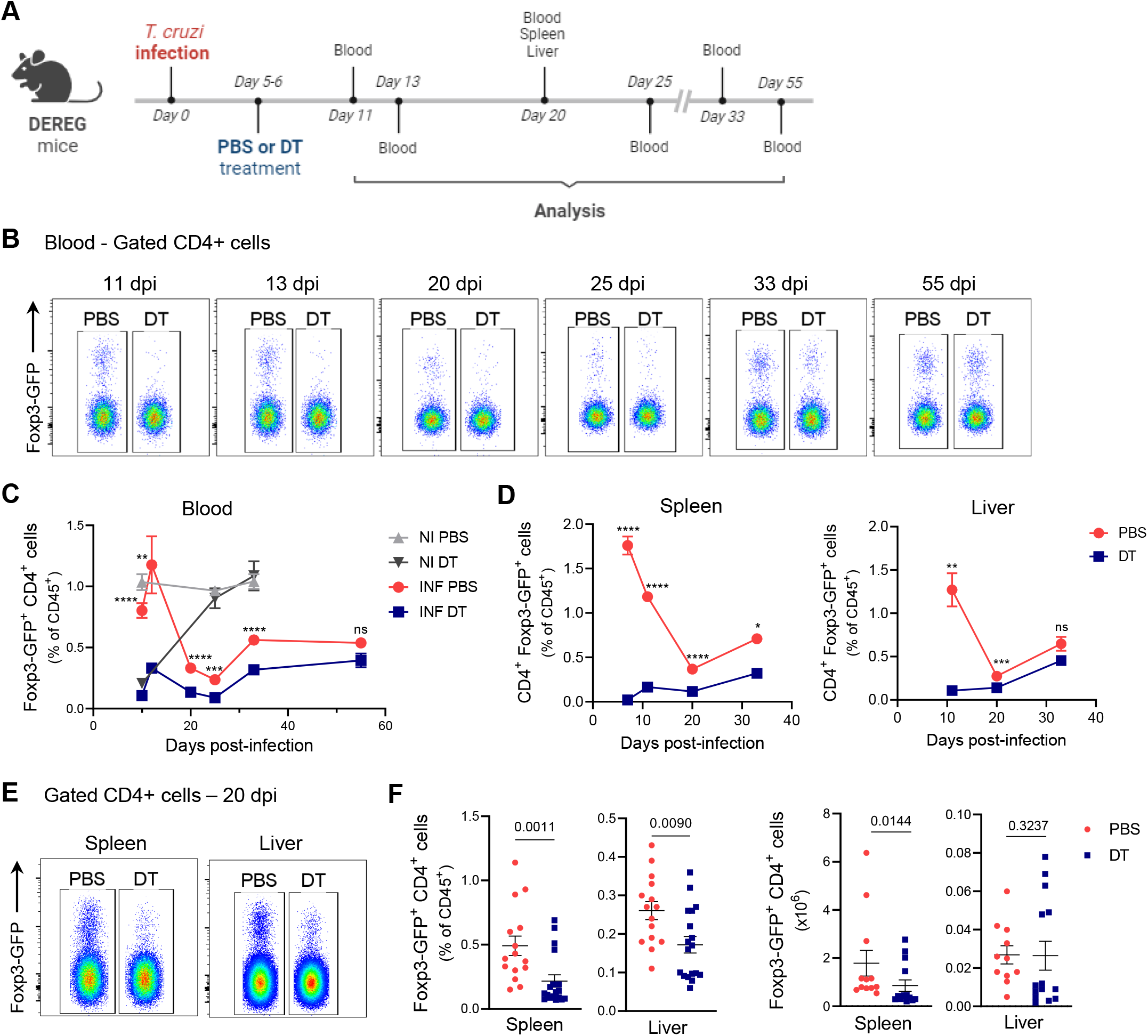
DT injection efficiently depletes Treg cells in DEREG mice during acute *T. cruzi* infection. **A)** Experimental scheme illustrating DT treatment (created with BioRender.com). **B-C)** Representative flow cytometry dot plots depicting Foxp3-GFP expression (B) and Treg cell frequencies (C) in the blood of PBS or DT-treated *T. cruzi*-infected (INF) DEREG mice at different days post-infection (dpi) and non-infected (NI) controls. **D)** Kinetics analysis of Treg cell frequencies in spleen and liver of PBS or DT-treated *T. cruzi*-infected DEREG mice. **E-F)** Representative flow cytometry dot plots showing Foxp3-GFP expression (E) and Treg cells frequencies and absolute numbers (F) in the spleen and liver of PBS or DT-treated DEREG mice at day 20 pi. All data are presented as mean ± SEM. In (C) and (D) data were collected from 1-3 independent experiments. A total of 2-18 mice per group were included (minimum 2 mice/group/experiment). In (F) data were collected from 4 independent experiments, with each symbol representing one individual mouse. Statistical significance was determined by Unpaired t test or Mann Whitney test, according to data distribution. P values for pairwise comparisons are indicated in the graphs. Statistical analysis in (C) represents pairwise comparisons between INF PBS and INF DT groups. * P ≤ 0.05, ** P ≤ 0.01, *** P ≤ 0.001, **** P ≤ 0.0001 and ns = not significant.

However, in DT-treated infected animals, Treg cell frequencies remained significantly decreased until day 33 pi compared to Treg-sufficient infected controls. Thus, DT injection effectively intensified and prolonged the natural contraction of the Treg cell response that we and others have previously reported for *T. cruzi* infection (3, 13, 24). We further examined the spleen and the liver, a target tissue for *T. cruzi* that exhibits a substantial leukocyte infiltration during the acute phase of the infection (25–27). Importantly, Treg cell depletion was also observed in frequency in the spleen and liver of DEREG mice, as well as in absolute numbers in the spleen, including on day 20 pi when the strongest natural contraction of the Treg cell response was detected (Figure 1, D, E and F).

Consistent with our previous report [13], we found that Treg-depleted infected animals showed reduced levels of blood parasitemia (Figure 2A) and tissue parasite burden in *T. cruzi* target tissues, such as heart and liver, compared to their control counterparts at day 20 pi (Figure 2B), which coincides with the peak of the CD8+ T cell response in our model (23, 28). However, no differences in parasite load were observed in the spleen (Figure 2B). Considering the critical role of CD8+ T cells in controlling *T. cruzi* replication (29), we quantified CD8+ T cells specific for the immunodominant epitope TSKB20 (*T. cruzi* trans-sialidase amino acids 569-576 –ANYKFTLV–) (30) in blood, spleen and liver (Figure 2C). Through kinetics studies in blood, we determined that Treg cell depletion significantly increased the frequencies of circulating TSKB20-specific CD8+ T cells from day 20 to day 33 pi in DT-treated animals compared to controls (Figure 2, C and D). Subsequently, the response progressively contracted, reaching frequencies similar to PBS-treated mice at day 55 pi. Similar results were observed in the spleen and liver, where higher relative and absolute numbers of parasite-specific CD8+ T cells were observed in Treg-depleted animals compared to Treg-sufficient mice at day 20 pi (Figure 2, E and F). Importantly, the observed effects on the *T. cruzi*-specific CD8+ T cell response and parasite levels were not attributable to potential toxic effects of DT itself but a consequence of Treg depletion, as they were not detected in DT-treated WT littermates (Supplemental Figure 1A). Additionally, Treg cell depletion had no effect on the plasmatic levels of biochemical markers of tissue damage and/or general health (Supplemental Figure 1B), except for a tendency towards a reduction in GOT activity, suggesting that a more robust effector response is not necessarily linked to increased acute tissue damage.

**Figure 2.**
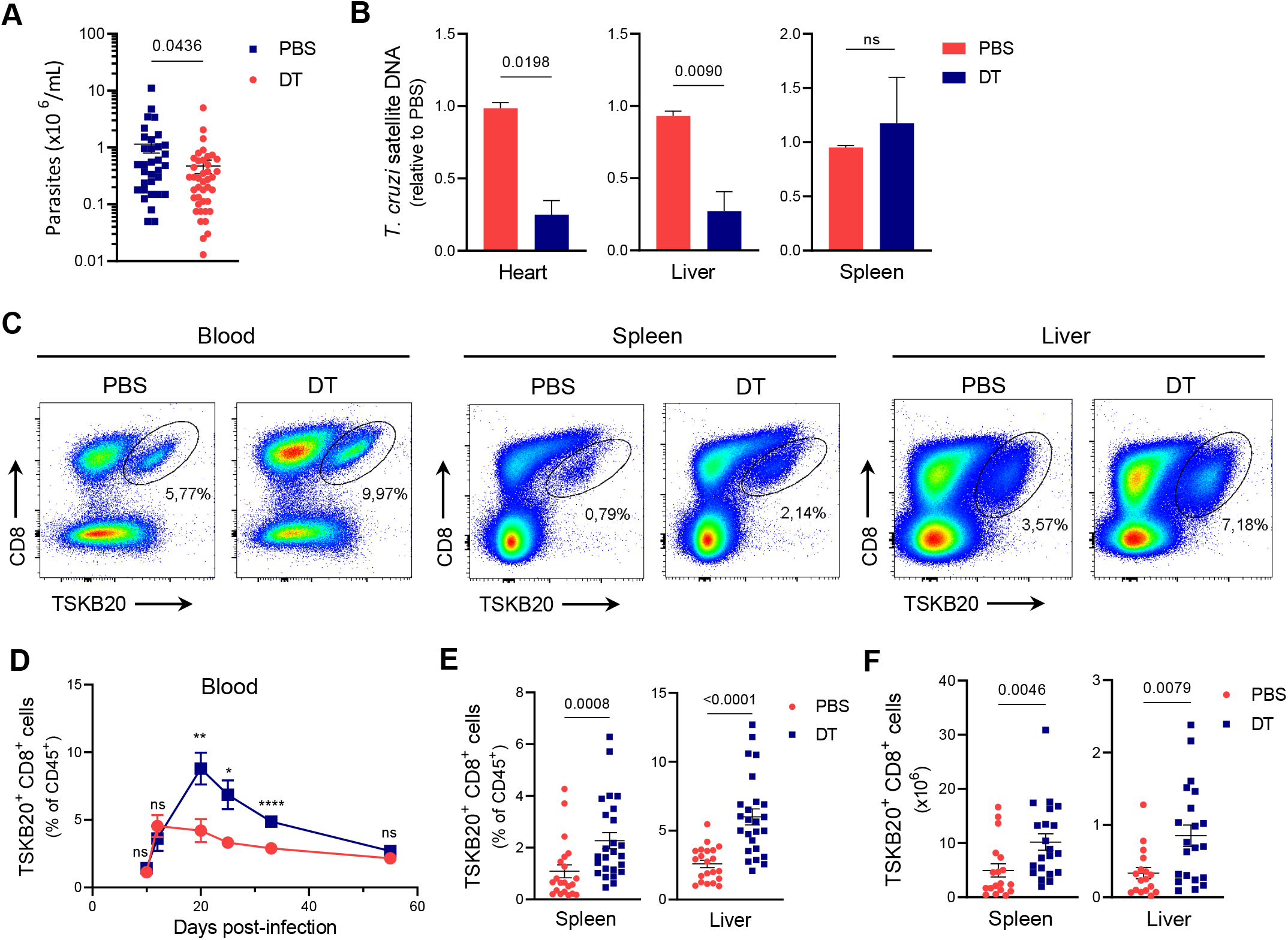
Treg cell depletion increases *T. cruzi*-specific CD8+ T cell expansion and improves parasite control during acute infection. **A)** Parasitemia from PBS or DT-treated DEREG mice at day 20 pi. **B)** Parasite load in heart, liver, and spleen of PBS or DT-treated DEREG mice at day 20 pi. **C-E)** Representative flow cytometry dot plots showing TSKB20-specific CD8+ T cell detection (C) and their frequency quantification (D, E) in blood, spleen and liver of PBS or DT-treated DEREG mice at day 20 pi (C, E) and different dpi (D). **F)** Absolute numbers of TSKB20-specific CD8+ T cells in spleen and liver according to (E). All data are presented as mean ± SEM. Data were collected from 1-2 independent experiments at most dpi, and 8 independent experiments at day 20 pi. In (B) data were pooled from 2-4 independent experiments. In (D) data correspond to 1-3 independent experiments. In (E-F) each symbol represents one individual mouse and data were collected from 4-5 independent experiments. In (A) to (F) a total of 4-42 mice per group were included (minimum 4 mice/group/experiment). Statistical significance was determined by Unpaired t test in (B) and Mann Whitney test in (A), (D), (E) and (F). P values for pairwise comparisons are indicated in the graphs. * P ≤ 0.05, ** P ≤ 0.01, *** P ≤ 0.001, **** P ≤ 0.0001 and ns = not significant.

Next, we studied whether the role of Treg cells in regulating the effector response was time-dependent during the acute phase of *T. cruzi* infection. To address this, we modified the DT injection schedule to days 11 and 12 pi, aiming to deplete Treg cells at the time point when the natural contraction of the Treg cell response begins and the anti-parasite T cell response is already detectable. Surprisingly, despite Treg cells remained depleted in blood and spleen until at least day 21 pi (Supplemental Figure 2, B and C), the delayed Treg cell depletion strategy had no impact on parasitemia levels or the frequencies of *T. cruzi*-specific CD8+ T cells in the blood (Supplemental Figures 2, D and E). Furthermore, no effects were observed in the frequencies of the anti-parasitic response at day 21 pi in the spleen of DT-treated animals, as compared to the control group (Supplemental Figure 2F).

Altogether, these results indicate that Treg cells play a prominent role during the initial stages of *T. cruzi* infection, likely influencing the priming and/or activation of the effector CD8+ T cell response. The suppressive function of Treg cells during the early phase of infection ultimately impacts the magnitude of the TSKB20-specific CD8+ T cell response and, consequently, the control of parasite replication.

### Early depletion of Treg cells promotes differentiation of parasite-specific CD8+ T cells into polyfunctional short lived effector cells

To examine the influence of Treg cell depletion on specific subsets of CD8+ T cells in the context of *T. cruzi* infection, we investigated the contribution of effector, memory and activated cells. Using flow cytometry, we identified short-lived effector cells (SLEC) and memory precursor effector cells (MPEC) based on CD44, KLRG-1 and CD127 expression, as illustrated in Supplemental Figure 3A (31, 32). Our analysis revealed that early Treg cell depletion increased the frequency of the SLEC subset within total and *T. cruzi*-specific CD8+ T cells from the spleen as well as the frequency and absolute numbers of SLEC within liver parasite-specific CD8+ T cells (Figure 3, A and B). In contrast, the MPEC subset did not show significant changes in frequency or absolute numbers in parasite-specific CD8+ T cells from the spleen and liver after DT injection (Figure 3C). Additionally, no significant differences in the frequencies of total or parasite-specific CD8+ T cells expressing the activation markers CD44 and CD25 were observed between DT-treated and PBS-treated mice (Supplemental Figure 3, C and D), indicating that the increase in the SLEC subset was the primary consequence of Treg depletion on CD8+ T cell responses at day 20 pi.

**Figure 3.**
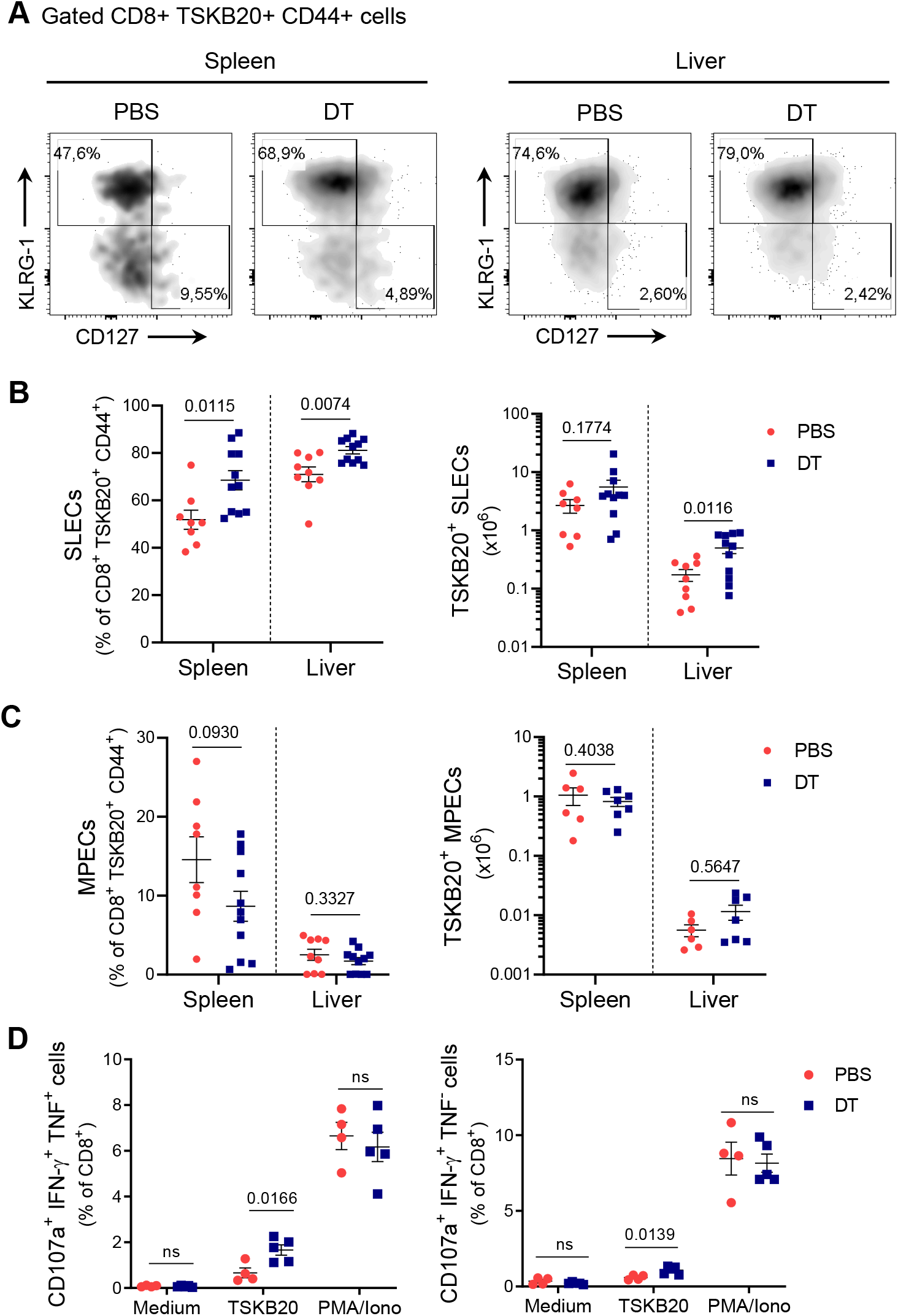
Early Treg cell depletion promotes parasite-specific CD8+ T cell differentiation into SLEC during *T. cruzi* infection. **A)** Representative flow cytometry density plots showing KLRG-1+ CD127-(SLEC) and KLRG-1-CD127+ (MPEC) subsets within CD44+ gated TSKB20-specific CD8+ T cells in the indicated organs obtained from PBS or DT-treated DEREG mice at day 20 pi. **B-C)** Frequencies (left) and absolute numbers (right) of SLEC (B) and MPEC (C) subsets within CD44+ gated TSKB20-specific CD8+ T cells of mice in (A). Data were collected from 2 independent experiments. **D)** Percentages of CD8+ T cells that produce IFN-γ and exhibit CD107a mobilization together (left) or not (right) with TNF production in the spleen of PBS or DT-treated DEREG mice at day 21 pi. Medium condition was used as a negative control, while PMA/Ionomycin (PMA/Iono) was used as a positive control for polyclonal CD8+ T cell stimulation. Similar results were obtained in 2 independent experiments. All data are presented as mean ± SEM. In (B), (C) and (D) each symbol represents one individual mouse. Statistical significance was determined by Unpaired t test or Mann Whitney test, according to data distribution. P values for pairwise comparisons are indicated in the graphs.

Based on our preceding results, we further investigated the functionality of parasite specific CD8+ T cells after DT treatment. To test this, we examined the degranulation ability by CD107a surface mobilization together with secretion of the effector cytokines IFN-γ and TNF upon *in vitro* specific and polyclonal stimulation (Supplemental Figure 4A). As shown in Figure 3D, CD8+ T cells from 21 day-infected Treg-depleted mice exhibited an increased proportion of polyfunctional cells that produce two and three effector mediators (degranulation plus IFN-γ production with or without TNF release) when stimulated with TSKB20 but not after polyclonal stimulation with PMA/Ionomycin. No significant changes were detected in other parasite-specific cells expressing one or two effector mediators (Supplemental Figure 4B). Altogether, our results indicate that early Treg cell depletion during *T. cruzi* infection enhances CD8+ T cell immunity by promoting the differentiation of effector cells with polyfunctional attributes.

### Effects of early Treg cell depletion on antigen-presenting cell populations and conventional CD4+ T cells

Next, we investigated whether other cell populations could be modulated by Treg cells at early stages of the infection and indirectly impact CD8+ T cell responses. Considering that only early depletion of Treg cells resulted in the increase of parasite-specific effector CD8+ T cells and improved control of parasite replication at day 20 pi, and taking into account that the TSKB20-specific CD8+ T cell response starts to arise around day 10 pi in our infection model (Figure 2D) (23), we hypothesized that Treg cells may play a role at early events of T cell priming during *T. cruzi* infection. To address this question, we evaluated the response of antigen-presenting cells (APCs) and innate immune cells the following day after DT treatment in the spleen of DT or PBS-treated and infected mice and non-infected controls, using multiparametric flow cytometry. As depicted in the Uniform Manifold Approximation and Projection (UMAP) visualization plots of Figure 4A, unsupervised analysis using the X-Shift clustering algorithm detected 10 clusters which were identified based on their expression of cell population markers (Supplemental Figure 5, A and B). Among these clusters, we observed six that displayed APC features, as evidenced by their expression patterns of MHC class I and class II molecules, along with activation costimulatory markers such as CD80, CD86 (33, 34), and CD24 (35) (Figure 4B). These clusters were designated as dendritic cells subset 1 (cluster 2), B cells (cluster 3), neutrophils (cluster 4), dendritic cells subset 2 (cluster 5), monocytes (cluster 7), and NKT cells (cluster 10).

**Figure 4.**
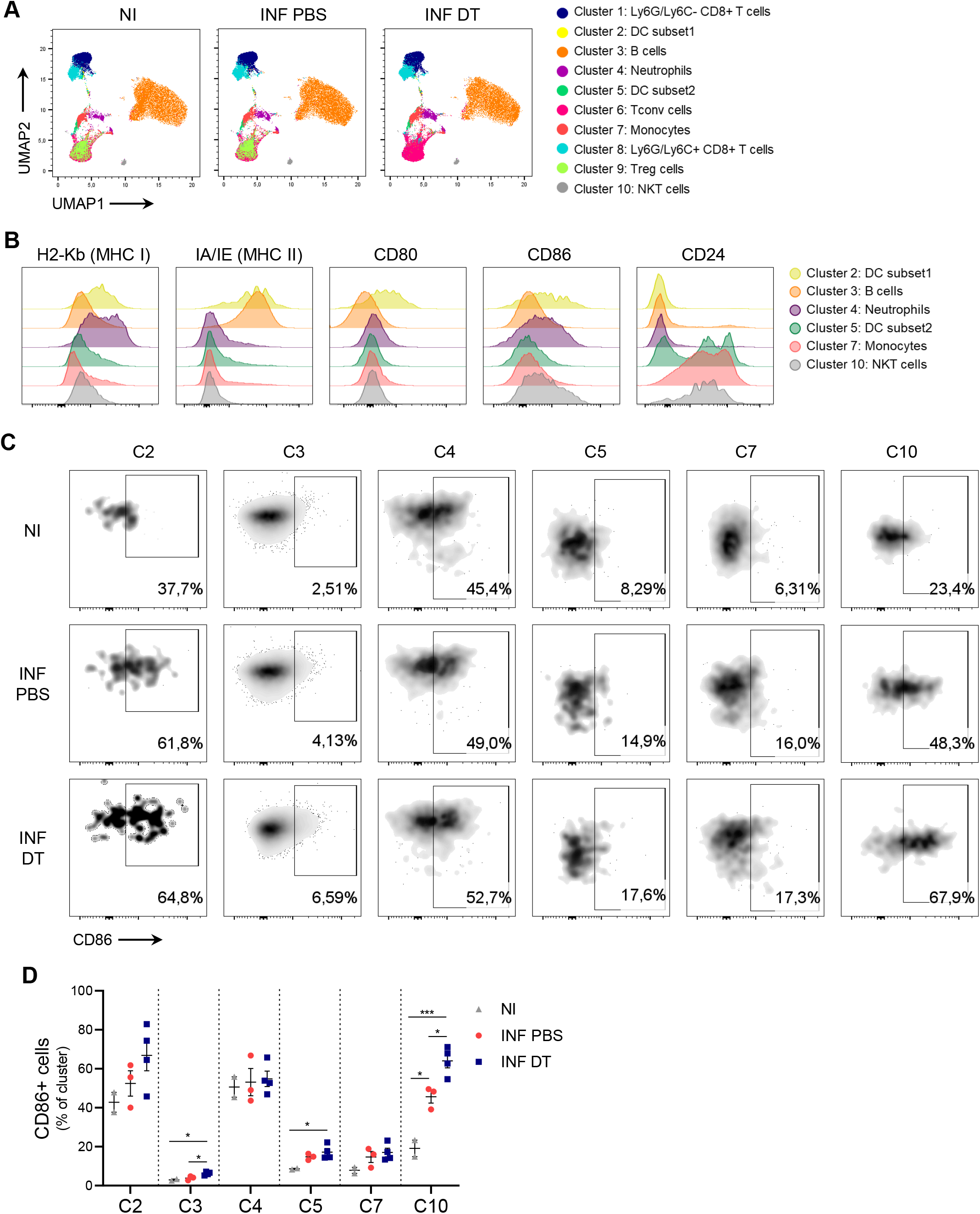
Treg cell depletion induces modest effects on APC populations and innate cells. **A)** UMAP visualization of flow cytometry data from the spleen of PBS or DT-treated DEREG mice at day 7 pi and non-infected controls. **B)** Histograms showing the expression of different APC and innate cell activation markers in selected cell clusters. Samples from the three experimental groups (NI, INF PBS and INF DT) were pooled together. **C)** Representative flow cytometry plots showing CD86+ cells in the indicated cell clusters as defined in (A). **D)** Frequency of CD86+ cells in the indicated cell clusters as defined in (A). Data are presented as mean ± SEM. Each symbol represents one individual mouse. Statistical significance was determined by one-way ANOVA followed by Tukey’s multiple comparison test. Similar results were obtained in 3 independent experiments. * P ≤ 0.05, *** P ≤ 0.001.

To assess the impact of Treg cell depletion on the frequency and phenotype of APCs and innate immune cells, we conducted a supervised analysis of our cytometric data. None of the six identified clusters showed significant changes in frequency between non-infected (NI) samples, Treg-depleted and non-depleted infected samples at the studied time point (Supplemental Figure 5C). However, we found that Treg cell depletion led to increased frequencies of CD86+ cells within the large cluster of B cells (C3) and, surprisingly, in NKT cells (C10), which showed an unexpected great proportion of cells expressing this co-stimulatory molecule (Figure 4, C and D). A similar trend was also observed in one DC cluster (C2). Interestingly, DT-treated mice showed increased frequency of CD86+ cells compared to non-infected controls in the clusters of B cells (C3), DC subset 2 (C5) and NKT cells (C10). No significant differences were observed in the frequency of positive cells for the remaining markers studied across the experimental groups within any of the identified clusters (Supplemental Figure 5D). Taken together, these data point to subtle effects of Treg cells on APC and innate immune cell populations at early stages following *T. cruzi* infection.

A previous study has demonstrated that CD4+ T cell help is required to mount a full-sized TSKB20-specific CD8+ T cell response to *T. cruzi* infection (5). Since Foxp3-CD4+ T (Tconv) cells are potential targets for Treg cell suppression (36–38), we investigated the impact of Treg cell depletion on this effector cell population. At day 20 pi, animals that received PBS or DT on days 5 and 6 pi showed similar frequencies of Tconv cells in blood, spleen, and liver (Supplemental Figure 6A). However, we observed a significant increase in the percentage of this cell subset at day 11 pi in those tissues from Treg-depleted mice, along with a corresponding increase in the absolute numbers of Tconv in the liver (Figure 5, A and 5). Analysis of subset distribution based on differentiation status revealed that effector Tconv cells were increased in the spleen and liver of DT-injected mice at day 11 pi, at the expense of a reduction in the frequency of naïve or naïve and memory Tconv cell subsets, respectively (Figure 5, C and D). Moreover, an increased frequency of Tconv cells expressing the T cell activation markers KLRG-1 and CD25 was observed in the spleen, but not in the liver, of Treg-depleted animals at day 11 pi (Figure 5, E and F, and Supplemental Figure 6C). These results suggest that Treg cells influence helper responses by modulating Tconv cell activation state and numbers at early stages of *T. cruzi* infection coincident with the initiation of the parasite-specific CD8+ T cell response.

**Figure 5.**
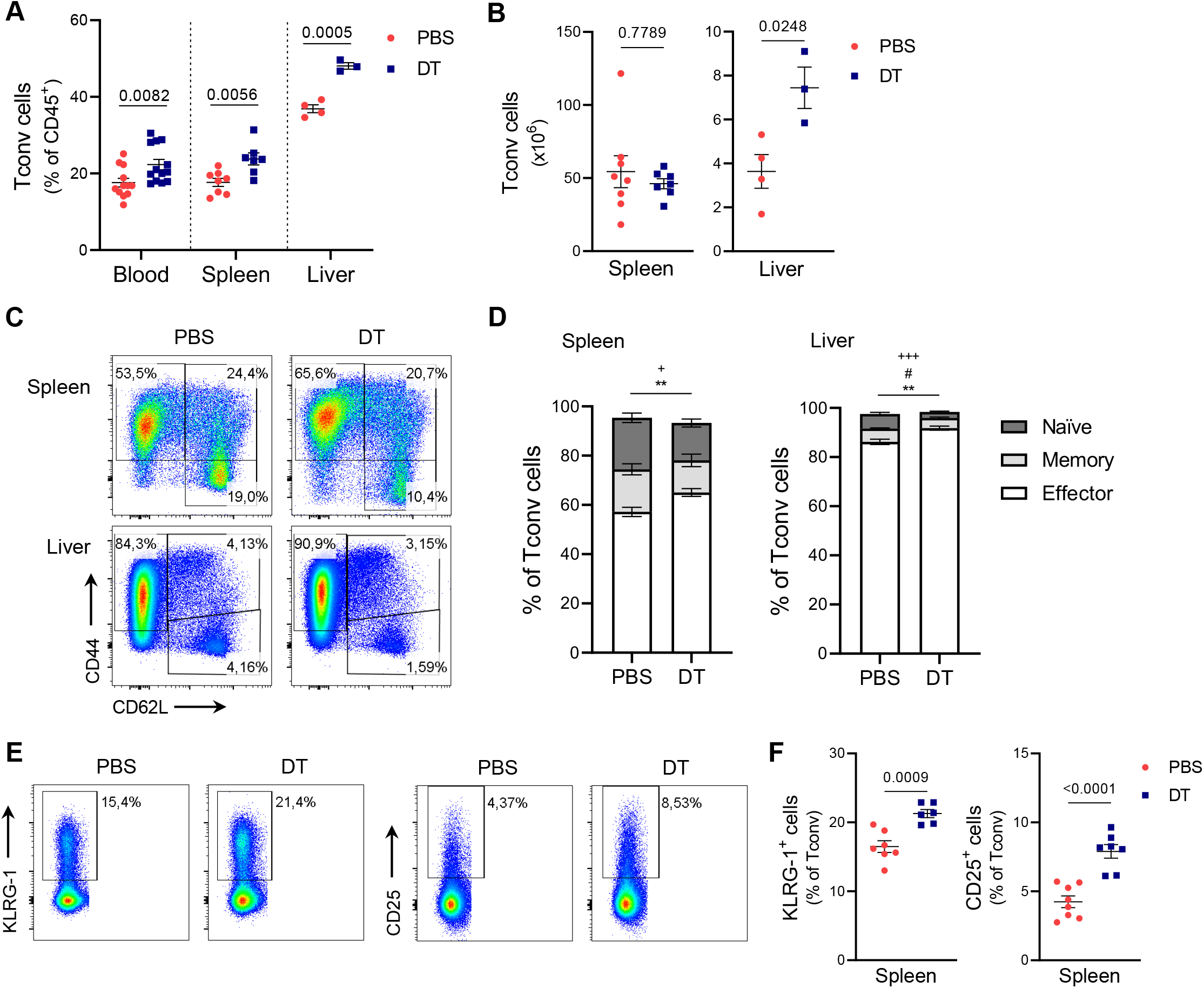
Treg cell depletion promotes the expansion and activation of Tconv cells in *T. cruzi* target organs. **A-B)** Frequencies (A) and absolute numbers (B) of Tconv cells in blood, spleen, and liver from PBS or DT-treated DEREG mice at day 11 pi. **C-D)** Representative flow cytometry plots (C) and frequency (D) of CD44-CD62L+ (naïve), CD44+ CD62L+ (memory) and CD44+ CD62L-(effector) Tconv cell subsets in the spleen and liver from mice in (A). **E-F)** Representative flow cytometry plots (E) and frequency (F) of KLRG-1+ and CD25+ Tconv cells in the spleen from mice in (A). All data are presented as mean ± SEM. Each symbol represents one individual mouse. Data in (A), (B), (D) and (F) were pooled from 1-2 independent experiments. Statistical significance was determined by Unpaired t test or Mann Whitney test, according to data distribution. P values for pairwise comparisons are indicated in the graphs. In (D), P values PBS vs DT: + P≤ 0.05 and +++ P ≤ 0.001, naïve Tconv cells; # P<0.05, memory Tconv cells; ** P ≤ 0.01, effector T conv cells.

### TSKB20-specific CD8+ T cell suppression is mediated by CD39 expression on Treg cells

In order to gain mechanistic insight into how Treg cells impact on the magnitude of the parasite-specific CD8+ T cell response, we then focused on suppressive molecules upregulated by Treg cells after *T. cruzi* infection. It is long known that Treg cells constitutively express key molecules associated with their suppressive function, including CTLA-4 and CD25 in addition to Foxp3 (39). In line with our previous findings that highlighted CD25, CTLA-4, and CD39 among the most upregulated markers in Treg cells during the peak of effector CD8+ T cell immunity (13), we sought to determine whether these molecules were also upregulated at earlier time points. Therefore, we analyzed their expression by flow cytometry at day 7 pi. Interestingly, while the frequency of CD25+ and CTLA-4+ Treg cells showed no differences between infected and non-infected mice at this time point, there was a significant increase in the proportion of CD39+ Treg cells in the spleen of *T. cruzi-*infected animals compared to controls, and importantly, of Treg cells expressing high levels of CD39 (Figure 6, A and B).

**Figure 6.**
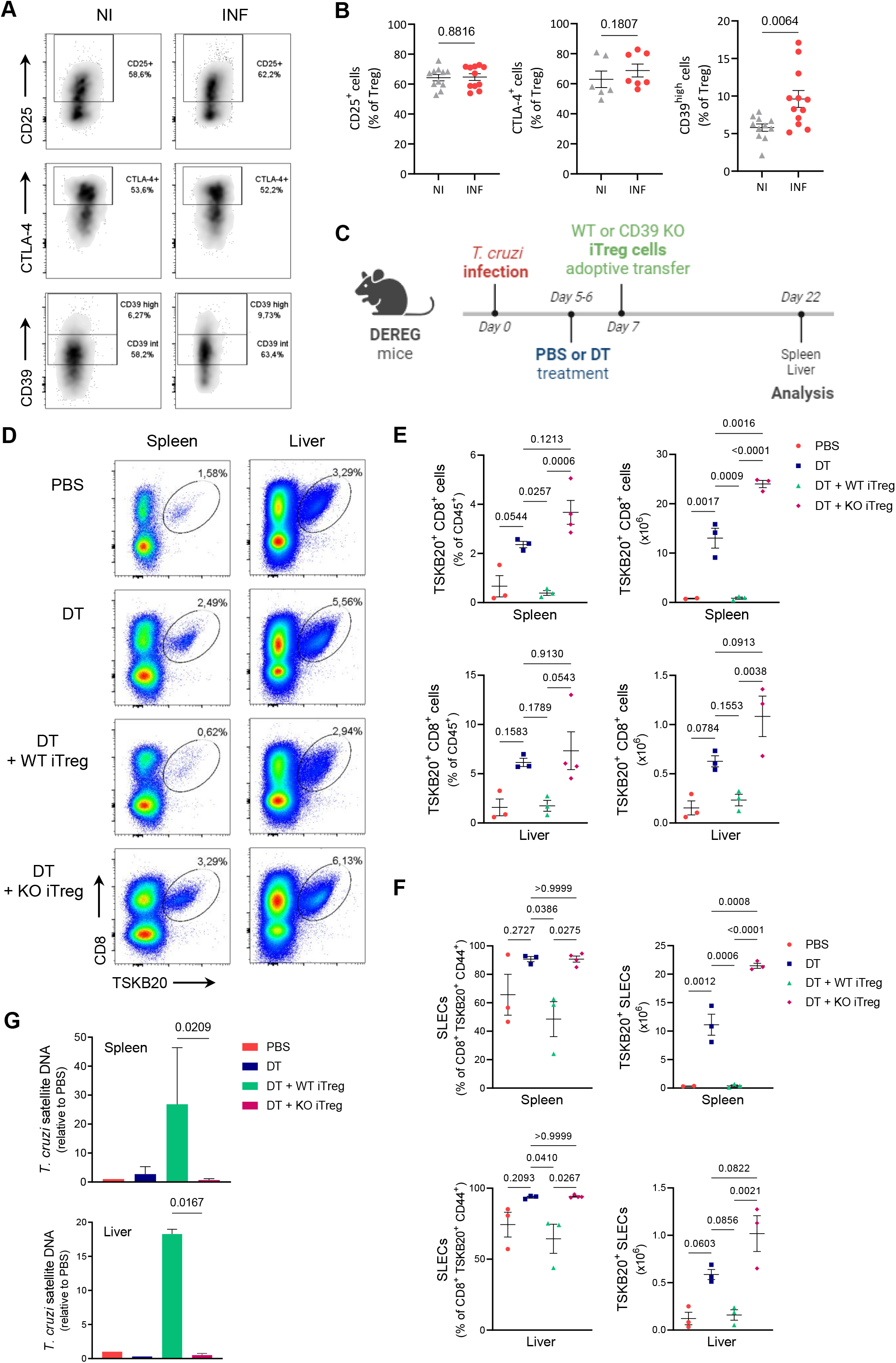
CD39 expression on Treg cells mediates TSKB20-specific CD8+ T cell suppression and parasite outgrowth. **A-B)** Representative flow cytometry plots (A) and frequency (B) of Treg cells expressing CD25, CTLA-4 and CD39 in the spleen of DEREG mice at day 7 pi (INF) and non-infected controls (NI). Data were collected from 2-3 independent experiments. **C)** Overview of iTreg cell adoptive transfer experimental design (created with BioRender.com). **D-E)** Representative flow cytometry plots (D) and frequency and absolute numbers (E) of TSKB20+ CD8+ T cells in the spleen and liver of mice treated as indicated in (C) at day 22 pi. **F)** Frequencies and absolute numbers of the SLEC subset within CD44+ gated TSKB20-specific CD8+ T cells from mice treated as indicated in (C). **G)** Parasite load in the spleen and liver from mice treated as indicated in (C). Results similar to (E) and (F) were obtained in 2 independent experiments. All data are presented as mean ± SEM. In (B), (E), and (F) each symbol represents one individual mouse. Statistical significance was determined by Unpaired t test or Mann Whitney test in (B), by one-way ANOVA followed by Tukey’s multiple comparisons test in (E) and (F), and by paired t test in (G). Values for pairwise comparisons are indicated in the graphs.

We then performed adoptive transfer experiments to deeply evaluate if the expression of CD39 by Treg cells is necessary for suppressing the TSKB20-specific CD8+ T cell response after *T. cruzi* infection. As illustrated in Figure 6C, DEREG mice were infected and Treg cell depletion was carried out as usual by DT administration at days 5 and 6 pi. The following day, a group of DT-treated mice received *in vitro* differentiated Treg (iTreg) cells from CD4+ splenocytes of non-infected WT or CD39 KO animals. As expected, Treg cell depletion led to an increased TSKB20-specific CD8+ T cell response at day 22 pi in the spleen and liver (Figure 6, D and E). Notably, depleted animals that were injected with WT iTreg cells reversed the effect of DT treatment, as demonstrated by a reduction in the parasite-specific CD8+ T cell response to levels similar to those of PBS-injected (Treg-sufficient) controls. In contrast, injection of CD39 KO iTreg cells in depleted mice resulted in a robust *T. cruzi*-specific CD8+ T cell response comparable to or higher than DT-treated animals, and notably higher than Treg-depleted mice that received WT iTreg cells, in both the spleen and liver (Figure 6, D and E). Consistent with our previous findings, the majority of TSKB20-specific CD8+ T cells displayed a SLEC phenotype at day 22 pi in the spleen and liver of Treg-depleted mice (Figure 6F). Interestingly, the relative and absolute numbers of SLEC TSKB20-specific CD8+ T cells significantly decreased when Treg-depleted animals received WT iTreg cells, but not when they were transferred with CD39 KO iTreg cells. Importantly, *T. cruzi* load in spleen and liver was significantly reduced when Treg-depleted mice were transferred with CD39 KO iTreg cells compared to mice that received WT iTreg cells (Figure 6G).

Collectively, these findings provide compelling evidence for the involvement of CD39-expressing Treg cells in modulating the magnitude of the effector *T. cruzi*-specific CD8+ T cell response and the control of parasite replication during the acute phase of infection. Importantly, this regulatory function was not compensated by other suppressive molecules expressed in Foxp3+ regulatory T cells in the context of *T. cruzi* infection.

## Discussion

Regulatory T cells play a pivotal role in infections by maintaining an effective balance between efficient immune responses for pathogen clearance and excessive immune reactivity that can lead to tissue damage. While the accumulation of Treg cells in the chronic phase has been well-documented for various infections, the contribution to immunity and disease outcome during the acute phase appears more highly context-dependent (40).

In this study, we demonstrate the crucial role of Treg cells in modulating the anti-parasite immune response and controlling pathogen replication during the acute phase of experimental *T. cruzi* infection. This work complements our previous study on the Treg cell response (13). Here, we have elucidated the timing and mechanisms through which activated Treg cells, despite a relative decrease in numbers, exert a suppressive function on effector CD8+ T cell immunity during acute *T. cruzi* infection. These results differ from previous observations where discrepancies were reported regarding the role of Treg cells in this parasitic infection, as some studies described beneficial impacts (24, 41), while others suggested limited (42, 43) or deleterious effects (44). However, none of these studies have thoroughly investigated the phenotypical and functional characteristics of the Treg cell response, and more importantly, all of them targeted these cells by non-specific approaches.

Taking advantage of a diphtheria toxin (DT) model for Treg cell depletion, we found that removal of endogenous Treg cells shortly after infection, but not at later stages, improved parasite control. We consider this effect is secondary to enhancement of TSKB20-specific CD8+ T cell immunity without causing significant tissue damage at the peak of the CD8+ T cell response. These results align with a previous study that reported increased numbers of CD8+ T cells restricted to the VNHRFTLV epitope of Amastigote Surface Protein-2 from *T. cruzi* when Treg cells were ablated (14). Overall, these observations highlight the role of Treg cells in suppressing optimal CD8+ T cell-mediated immunity against *T. cruzi* and indicate that Treg cells play highly relevant functions in this parasitic infection.

Following activation by antigen encounter, CD8+ T cells undergo expansion and differentiation into effector cells, which can be categorized into two distinct subsets: highly functional short-lived effector cells (SLEC) that will undergo apoptosis after antigen clearance, and memory-precursors (MPEC) that are able to continue the differentiation program into long-lived memory cells. In the current study, we provide evidence that Treg cells preferentially suppress the magnitude of the SLEC subset among parasite-specific CD8+ T cells, while having no impact on the generation of parasite-specific MPECs. Our results contrast with previous observations where Treg cells either promote (45–48) or restrict (49) the development of the CD8+ T cell memory response in viral or bacterial infections and immunization models. These discrepancies might be attributed to variations in the inflammatory milieu of each setting, considering the essential role of cytokines in directing CD8+ T cell differentiation programs, including IL-2 and IL-10 that mediate Treg cell function in those studies.

Multiple regulatory mechanisms have been described for Treg cells, such as the secretion of suppressive cytokines and cytolytic molecules, expression of inhibitory receptors and metabolic disruption of effector cells. However, there is still limited understanding of the precise mechanisms employed by Treg cells to mediate their suppressive activity in different inflammatory settings. Acquiring this knowledge will enable the modulation of inflammation outcomes associated with infections by therapeutic approaches targeting Treg cells. In recent years, purinergic suppressive pathways have gained significance. Treg cells have been shown to control inflammation in experimental models and in clinical studies, mediated, at least in part, through an adenosine-dependent manner via the regulated expression of CD39 and CD73 (17, 50–54).

In particular, it has been reported that human and mouse Treg cells expressing CD39 suppress Tconv and CD8+ T cell proliferation or cytokine production more efficiently *in vitro,* when compared to Treg cells lacking CD39 expression (55, 56). Additionally, CD39-expressing Treg cells exhibit stronger *in vitro* suppressive effects than CD39-Treg cells or upon CD39 inhibition (57–59). In our previous study, we reported that Treg cells upregulate CD39 expression along with a wide range of suppressive markers and regulatory molecules during the acute phase of *T. cruzi* infection (13).

In the current study, we show that *T. cruzi* infection soon increases the proportion of CD39-expressing Treg cells post-infection. Moreover, by adoptive transfer experiments, we can demonstrate that CD39 expression on Treg cells is necessary for controlling parasite-specific CD8+ T cells responses; independent of other regulatory mechanisms. To our knowledge, this is the first description of *in vivo* T cell suppression mediated by CD39+ Treg cells in the context of this model of acute *T. cruzi* infection. Our findings implicate Treg cells (and other cells) expressing CD39 as a possible pharmacological target for interventions aimed at modulating the magnitude of the effector anti-parasite response in *T. cruzi* infection. Remarkably, a recent report showed that patients with chronic chagasic cardiomyopathy exhibit expansion of activated effector CD4+ T cell subsets, which correlate with a decreased frequency of CD39+ Treg cells. These data suggest that suppressive mechanisms also operate in the context of the human infection (60). It would be interesting to investigate whether other components of the adenosine pathway are also involved in the control of immune responses in this parasitic infection. For instance, *T. cruzi* infection did not modify CD73 expression on Treg cells as shown in our previous study (13), suggesting that ATP conversion to adenosine might be dependent on other cells expressing CD73, which could act in concert with CD39+ Treg cells. Indeed, macrophages and neutrophils express higher levels of CD73, when compared to lymphocytes from the heart, visceral adipose tissue, and liver during experimental *T. cruzi* infection (61, 62).

Treg cell-mediated suppression of effector CD8+ T cell responses can occur through direct contact/proximity between the cells (63, 64), indicating a direct suppressive effect, or through the modulation of other immune cell populations that are important for CD8+ T cell functions (65), suggesting an indirect regulatory mechanism. In this study, we provide clear evidence that Treg cells not only impact CD8+ T cell immunity but also restrain Tconv cell responses during acute *T. cruzi* infection. Our observation that Treg cell depletion affects Tconv cell numbers and their activation phenotype prior to the regulation of parasite-specific CD8+ T cells suggests an indirect effect of Treg cells on parasite-specific CD8+ T cells through the modulation of Tconv cells. However, we cannot rule out the possibility of direct interactions between Treg cells and CD8+ T cells. CD4+ T cells have long been recognized for their role in providing help for CD8+ T cell priming, particularly in the generation of memory CD8+ T cell pools (66). The “licensing” model has been proposed as the main mechanism for CD4+ T cell help delivery, in which these cells are necessary to enhance the antigen-presenting and co-stimulatory capacities of DCs to induce robust effector CD8+ T cell responses (67). In agreement with our results, a study found that activation of CD4+ T helper cells preceded that of CD8+ T cells and that this process involved the action of different DC subsets in the context of HSV infection (68). In our system, APCs were only mildly affected by Treg cell depletion. Nevertheless, it would be interesting to evaluate the impact of early Treg cell elimination on APCs at later time points coinciding with the effect on the Tconv cell response in order to establish if a Treg-Tconv-APC axis is involved in the modulation of parasite-specific CD8+ T cells during *T. cruzi* infection. Furthermore, it has been reported that adenosine can downregulate CD86 expression on DCs (69, 70). Whether CD39 activity on Treg cells is responsible for the modulation of Tconv cell and APC responses needs to be further addressed in this parasitic infection.

NKT cells are a group of innate-like T cells that recognize lipids in the context of CD1d molecules. By responding very rapidly to TCR and/or cytokines they can link innate and adaptive immune responses and are important players in infectious diseases (71). The contribution of NKT cells to protection against *T. cruzi* remains controversial in the context of the experimental infection (72–77), while few studies have focused on this cell population in human Chagas disease. For instance, established chronic infection in asymptomatic patients can be accompanied by increased peripheral blood frequencies of both NKT cells and regulatory CD25+ T cells, and these values were inversely correlated to numbers of activated CD8+ T cells (78, 79). In this study, we found that *T. cruzi* infection augment the frequency of NKT cells expressing CD86 and that this effect was even increased after Treg cell depletion, potentially linked to CD8+ T cell priming during the acute phase. The implications of CD86 expression by NKT cells, the interaction between NKT cells and Treg cells, and their functional properties in the context of *T. cruzi* infection deserve further investigation.

In summary, the current study supports our previous hypothesis that weakened Treg cell responses during the acute phase of *T. cruzi* infection have beneficial elements for the host. This mechanism allows the generation of robust anti-parasite responses that prevent pathogen outgrowth without enhancing tissue damage. Further research is required to evaluate any sequelae of early Treg cell depletion. One potential outcome is that improved control of the parasite burden in the acute phase limits persistence of *T. cruzi* in target tissues during the chronic stage. Conversely, the reinvigorated effector response may prevail, resulting in immunopathology. Additionally, the restriction of Treg cells during the chronic phase should also be carefully examined, as this may trigger harmful inflammatory responses by leveraging chronic effector responses. In fact, previous studies of the human infection have linked impaired Treg cell immunity to more severe clinical forms of Chagas disease (78, 80, 81). Therefore, more detailed investigations are necessary to fully understand the role of Treg cells and CD39 expression in the transition from acute to chronic *T. cruzi* infection and the development of disease.

## Methods

### Mice

Male and female mice aged 8 to 12 weeks were used for experiments. C57BL/6 and BALB/c wild type mice were obtained from School of Veterinary, La Plata National University (La Plata, Argentina). DEREG (C57BL/6-Tg(Foxp3-DTR/EGFP)23.2Spar/Mmjax) reporter mice were purchased from The Jackson Laboratories (USA). CD39 deficient mice (82). All mouse strains were bred and housed at the animal facility of the Facultad de Ciencias Químicas, Universidad Nacional de Córdoba (FCQ-UNC). Mice were maintained on a 12-hour light/12-hour dark cycle with food and water *ad libitum*. Mice from different experimental groups were co-housed in the same cages. DEREG mice were bred as heterozygotes by breeding on a C57BL/6 background, and F1 generation was routinely tested for GFP expression.

### Parasites and experimental infection

Bloodstream trypomastigotes of the Tulahuén strain of *T. cruzi* were maintained in male BALB/c mice by serial passages every 10 - 11 days. For experimental infection, mice were intraperitoneally inoculated with 0.2ml PBS containing 5 × 10^3^ trypomastigotes. All infections were performed at similar hours of the day.

### Treg cell depletion

Mice were randomly divided into Treg cell depleted and control groups. DEREG mice were injected intraperitoneally with 25 ng of Diphtheria Toxin (DT; Calbiochem) per gram of body weight (25 ng/g) diluted in PBS. DT was administered in two consecutive days at the indicated time points. Control non-depleted mice received PBS injections. The following day, Treg cell ablation was confirmed in blood samples by flow cytometry.

### Parasite quantification

Parasitemia was assessed by counting the number of viable trypomastigotes in blood after lysis with a 0.87% ammonium chloride buffer. Abundance of *T. cruzi* satellite DNA in tissues was used to determine parasite burden. Genomic DNA was purified from 50 μg of tissue (heart, liver, and spleen) with TRIzol Reagent (Life Technologies) following manufacturer’s instructions. DNA samples from the same tissue and experimental group were pooled in each experiment. Satellite DNA from *T. cruzi* (GenBank AY520036) was quantified by real time PCR using specific Custom Taqman Gene Expression Assay (Applied Biosystems). Primers and probes sequences were previously described by Piron *et al*. (83). The samples were subjected to 45 PCR cycles in a thermocycler StepOnePlus Real-Time PCR System (Applied Biosystems). Abundance of satellite DNA from *T. cruzi* was normalized to the abundance of GAPDH (Taqman Rodent GAPDH Control Reagent, Applied Biosystems), quantified through the comparative ΔΔCT method and expressed as arbitrary units, as previously reported (13, 23, 28).

### Cell preparation

Blood was collected via cardiac puncture using heparin as anticoagulant. Spleens and livers were obtained and mechanically disaggregated through a tissue strainer to obtain cell suspensions in PBS 2% FBS. Liver infiltrating leukocytes were isolated by centrifugation at 600g for 25 minutes using a 35% and 70% bilayer Percoll (GE Healthcare) gradient. Erythrocytes were lysed for 3 min using an ammonium chloride-potassium phosphate buffer (ACK Lysing Buffer, Gibco). Cell numbers were counted in Turk’s solution using a Neubauer chamber.

### Biochemical determinations

Blood samples were centrifuged at 3000 rpm for 8 min and plasma was collected. Quantification of GOT, GPT, LDH and CPK activities was performed by UV kinetic method, CPK-MB activity by enzymatic method, and glucose concentration by enzymatic/colorimetric method at Biocon Laboratory (Córdoba, Argentina) using a Dimension RXL Siemens analyzer.

### Flow cytometry

For surface staining, cell suspensions were incubated with fluorochrome labeled-antibodies together with LIVE/DEAD Fixable Cell Dead Stain (ThermoFisher) in PBS 2% FBS for 20 min at 4°C. To identify *T. cruzi* specific CD8+ T cells, cell suspensions were incubated with an H-2Kb *T. cruzi* trans-sialidase amino acids 569-576 ANYKFTLV (TSKB20) APC- or BV412-labeled Tetramer (NIH Tetramer Core Facility) for 20 min at 4°C, in addition to the surface staining antibodies. To detect NKT cells, cell suspensions were incubated with PBS-57, an analogue of a-galactosylceramide (α-Gal-Cer) (84), complexed to CD1d APC-labeled Tetramers (NIH Tetramer Core Facility). Blood samples were directly incubated with the specified antibodies, and erythrocytes were lysed with a 0.87% NH4Cl buffer prior to acquisition.

For the detection of transcription factors, cells were initially stained for surface markers, washed, fixed, permeabilized and stained with Foxp3/Transcription Factor Staining Buffers (eBioscience) according to eBioscience One-step protocol for intracellular (nuclear) proteins.

For the UMAP analysis, the cytometry panel included the following anti-mouse monoclonal antibodies: PerCP-eFluor 710 anti-CD80 (B7-1) clone 16-10A1 (eBioscience), Super Bright 436 anti-MHC Class I (H-2kb) clone AF6-88.5.5.3 (eBioscience), Super Bright 600 anti-MHC Class II (I-A/I-E) clone M5/114.15.2 (eBioscience), Super Bright 645 anti-CD11b clone M1/70 (eBioscience), Super Bright 702 anti-Ly-6G/Ly-6C clone RB6-8C5 (eBioscience), Brilliant Violet 785 anti-CD86 clone GL-1 (Biolegend), Alexa Fluor 700 anti-CD45 clone 30-F11 (eBioscience), APC/Cyanine7 anti-F4/80 clone BM8 (Biolegend), PE anti-CD3e clone 145-2C11 (eBioscience), PE-eFluor 610 anti-CD24 clone M1/69 (eBioscience), PE-Cyanine5 anti-CD19 clone eBio1D3 (1D3) (eBioscience), PE-Cyanine5.5 anti-CD8a clone 53-6.7 (eBioscience), PE-Cyanine7 anti-CD11c clone N418 (eBioscience). The staining also included Tetramers to identify NKT cells and the LIVE/DEAD™ Fixable Aqua Dead Cell Stain Kit, for 405 nm excitation (Invitrogen).

The detailed list of antibodies used in all the experiments can be found in S1 Table. All samples were acquired using a FACSCanto II (BD Biosciences) or a LSRFortessa X-20 (BD Biosciences) flow cytometer, and data were analyzed with FlowJo software versions X.0.7 and 10.8.1.

### Determination of CD8+ T Cell effector function *in vitro*

Spleen cell suspensions were cultured for 5h in RPMI 1640 medium (Gibco) supplemented with 10% heat-inactivated FBS (Gibco), 2 mM glutamine (Gibco), 55 μM 2-ME (Gibco), and 40 μg/ml gentamicin. Cells were stimulated with 2.5 μM TSKB20 (ANYKFTLV) peptide (Genscript Inc.) or 50 ng/mL PMA plus 1 μg/ml ionomycin (Sigma-Aldrich) in the presence of Monensin and Brefeldin A (eBioscience). Culture medium was used as negative control. Anti-CD107a was included during the culture period. After surface staining, cells were fixed and permeabilized using Intracellular Fixation & Permeabilization Buffer Set (eBioscience) following manufacturer’s instructions. Stained cells were acquired on a FACSCanto II (BD Biosciences) or a LSRFortessa X-20 (BD Biosciences) flow cytometer as before. Antibodies specifications are detailed in S1 Table.

### Adoptive cell transfer

CD4+ T cells were isolated from pooled splenic cell suspensions by magnetic negative selection using EasySep™ Mouse CD4+ T Cell Isolation Kit (StemCell Technologies) according to manufacturer’s protocol. The isolated cells were further purified to obtain naïve CD4+ T cells (CD4+ CD25-CD44-) by cell sorting using a FACSAria II (BD Biosciences) instrument (see S1 Table for antibodies specifications). The sorted naïve CD4+ T cells were then incubated with a Treg cell differentiation cocktail in 96-well cell culture plates. For this, 2 x 10^5^ cells were stimulated with plate-bound anti-CD3 and anti-CD28 antibodies (eBioscience) at concentrations of 2 and 1 μg/ml, respectively, in the presence of 20 ng/mL rmIL-2 (Biolegend), 5 ng/mL m/h rTGF-β (eBioscience) and 13.3 nM *all trans*-Retinoic Acid (Sigma) diluted in supplemented RPMI 1640 medium (Gibco). On day 4, cells were harvested and Foxp3 expression was confirmed by intracellular staining. One million *in vitro* differentiated Treg cells were subsequently injected intravenously into the retro-orbital sinus of DEREG recipient mice infected with *T. cruzi* and previously treated with DT. Non-transferred and PBS-treated mice were included as controls.

### AI Language Model Assistance

We used ChatGPT (developed by OpenAI) to assist in refining the written content of this study. ChatGPT provided suggestions and corrections based on the input provided by the user, enhancing the clarity and grammar of the text. ChatGPT output was critically revised by the user to ensure it conveys the desired message.

### Statistics

Descriptive statistics were calculated for each experimental group. The normality of data distribution was assessed using Shapiro-Wilk normality test. Statistical significance of mean value comparisons was determined using two-tailed t-test or One-way ANOVA for normally distributed data, and two-tailed Mann Whitney test or Kruskal-Wallis test for non-normally distributed data, as appropriate. P values ≤ 0.05 were considered statistically significant. Outliers were identified using the ROUT method. GraphPad Prism 9.0 software was used for statistical analyses and graph creation. Data are presented as mean ± SEM. Sample size for each experiment is indicated in the figure legends, while the number of animals of each experimental group is shown in the scatter dot plots, unless stated otherwise.

### Study Approval

All animal procedures were conducted in compliance with the ethical standards set by the Institutional Animal Care and Use Committee of FCQ-UNC, and were approved under protocol numbers RD-731-2018 and RD-2134-2022.

## Author contributions

CLAF designed and conducted the experiments, acquired and analyzed data, and wrote the manuscript. SB and CR conducted experiments, contributed to investigation and methodology, and reviewed the manuscript. SCR provided resources and reviewed the manuscript. AG and CLM provided funding, contributed to investigation and methodology, and reviewed the manuscript. EVAR conceived and supervised the study, designed the experiments, provided funding and wrote the manuscript.

## Acknowledgments

We thank MP Abadie, MP Crespo, E Zacca, V Blanco, D Lutti, C Noriega, FA Frontera, SR Oms, RE Villarreal, G Furlán, NM Maldonado, A Romero and L Reyna (Centro de Investigaciones en Bioquímica Clínica e Inmunología) for their excellent technical assistance.

We acknowledge the NIH Tetramer Core Facility for provision of APC and BV421-labeled TSKB20 tetramers and APC-labeled PBS-57 complexed to CD1d tetramers.

This work was supported by: Agencia Nacional de la Investigación, el Desarrollo Tecnológico y la Innovación under grants PICT 2018-01791 and PICT 2020-0487, and the National Institute of Allergy and Infectious Diseases of the National Institutes of Health under Award Number R01AI169482. The content is solely the responsibility of the authors and does not necessarily represent the official views of the funding agencies.

## Notes

### Competing Interest Statement

The authors have declared no competing interest.

