## Supplemental Figures for "CD39 expression by regulatory T cells drives CD8+ T cell suppression during experimental *Trypanosoma cruzi* infection"

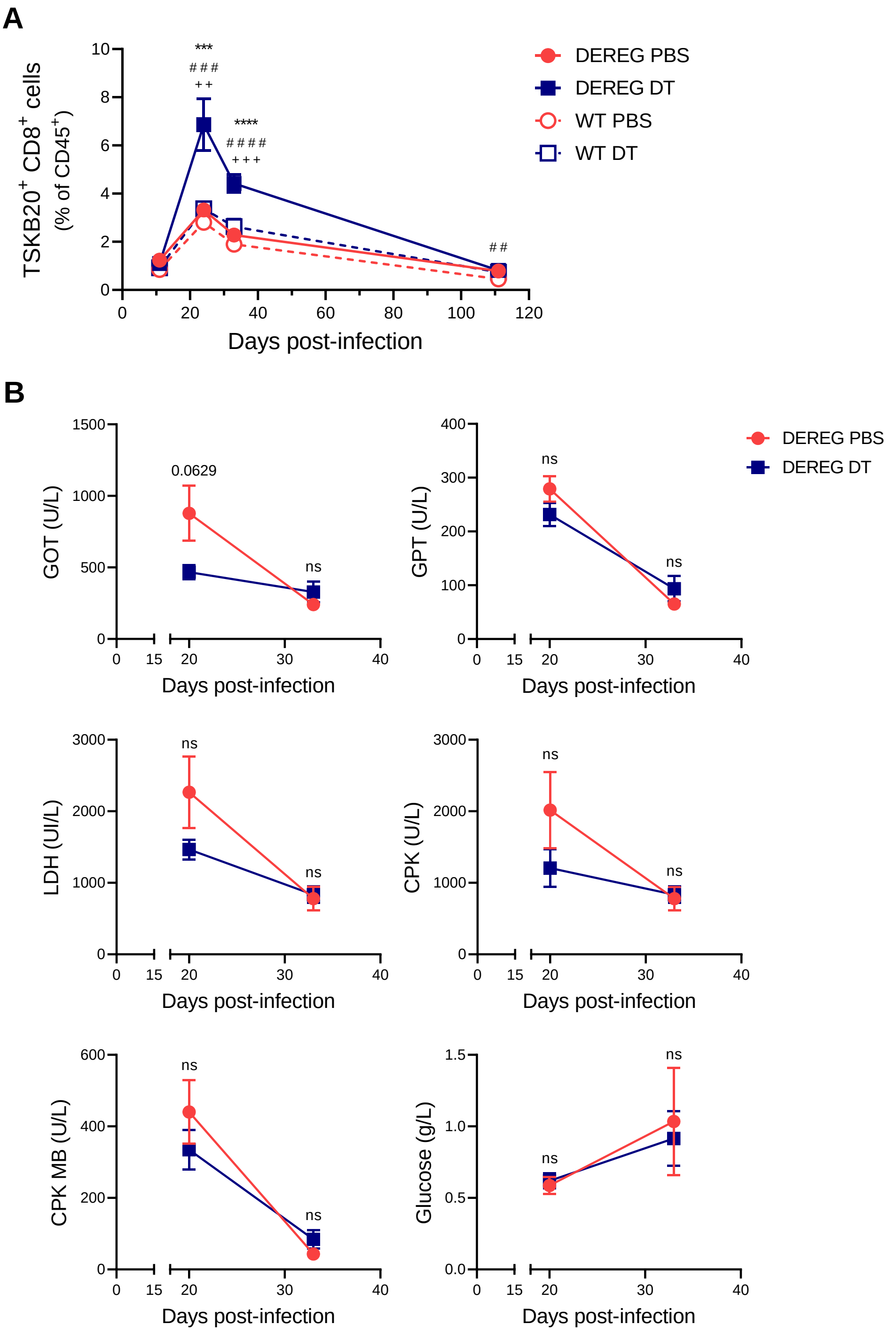


**Supplemental Figure 1. DT treatment controls. A)** TSKB20-specific CD8+ T cell frequencies in blood from PBS or DT-treated *T. cruzi* infected DEREG and WT littermate mice at different dpi. Data are presented as mean ± SEM. Statistical significance was determined by one-way ANOVA followed by Dunnett's multiple comparisons test. * DEREG DT vs DEREG PBS, # DEREG DT vs WT PBS, + DEREG DT vs WT DT. Similar results were obtained in 2 independent experiments. **B)** Treg cell depletion effect on tissue damage markers: activities of glutamate-oxalacetic transaminase (GOT), glutamate-pyruvate transaminase (GPT), lactate dehydrogenase (LDH), creatine phosphokinase (CPK), and creatine phosphokinase of muscle and brain (CPK MB), as well as Glucose concentration in plasma of PBS or DT-treated DEREG mice at days 20 and 33 pi. Data were collected from 1-2 independent experiments. Data are presented as mean ± SEM. Statistical significance was determined by Unpaired t test for GOT, LDH, CPK, and CPK MB activities and Glucose concentration, and by Mann Whitney test for GPT activity, according to data distribution. P values for pairwise comparisons at day 20 pi are indicated in the graphs.

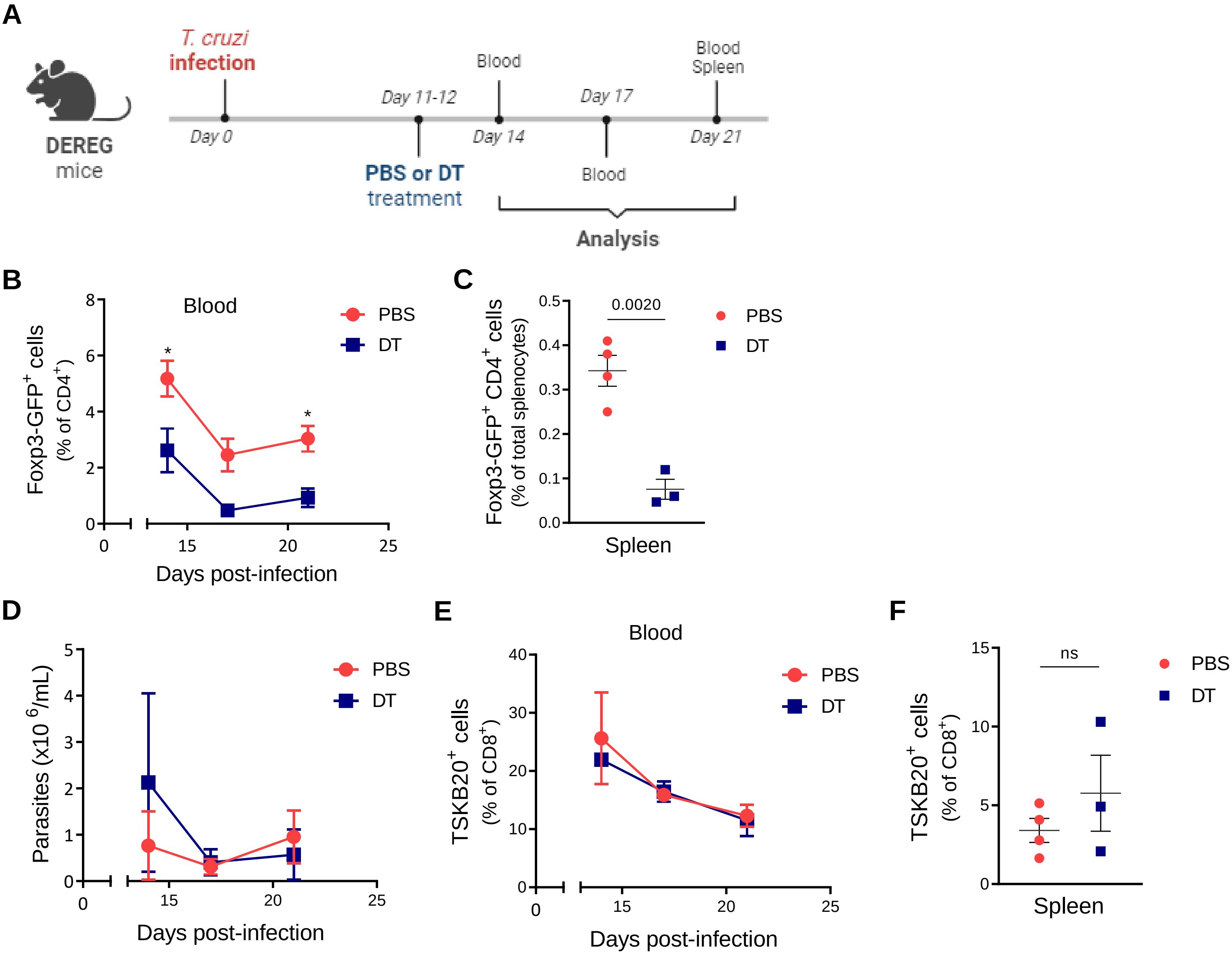


**Supplemental Figure 2. Depletion of Treg cells on days 11 and 12 pi had no impact on parasitemia levels or parasite-specific CD8+ T cell numbers. A)** Experimental scheme for DT treatment (created with BioRender.com). **B-C)** Treg cell frequencies in blood at different dpi (B) and in spleen at day 21 pi (C) from *T. cruzi*-infected DEREG mice treated with PBS or DT on days 11 and 12 pi. **D)** Parasitemia levels from mice in (A). **E-F)** TSKB20-specific CD8+ T cell frequencies in blood at different dpi (E) and in spleen at day 21 pi (F) of mice in (A). All data are presented as mean ± SEM. Data were collected from 1-2 independent experiments. Statistical significance was determined by Unpaired t test or Mann Whitney test, according to data distribution. P values for pairwise comparisons are indicated in the graphs. * P ≤ 0.05.

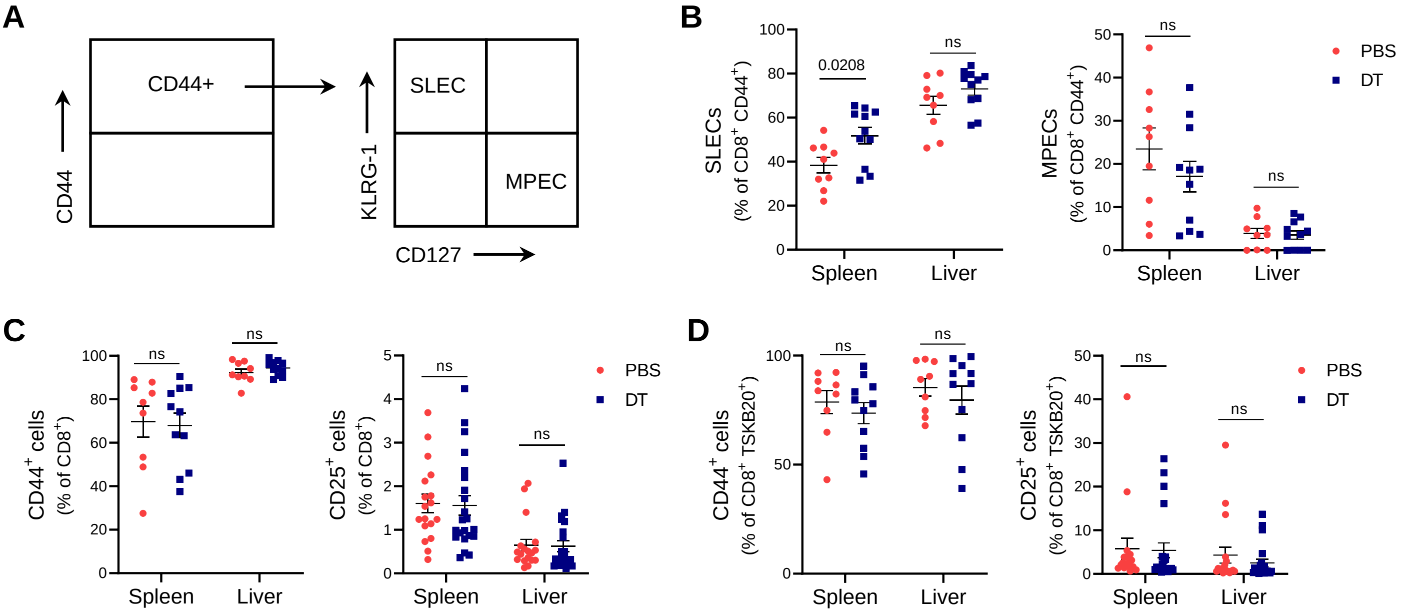


**Supplemental Figure 3. Effect of early Treg cell depletion on total and TSKB20-specific CD8+ T cell phenotype. A)** Gating strategy for evaluation of SLEC and MPEC subsets. **B)** Frequencies of SLEC (left) and MPEC (right) subsets within CD44+ gated CD8+ T cells from PBS or DT-treated DEREG mice at day 20 pi. **C-D)** Frequencies of cells expressing the indicated activation and exhaustion markers in total (C) and TSKB20-specific (D) CD8+ T cells in the spleen and liver of PBS or DT-treated DEREG mice at day 20 pi. Data were collected from 1-4 independent experiments and are presented as mean ± SEM. Each symbol represents one individual mouse. Statistical significance was determined by Unpaired t test or Mann Whitney test, according to data distribution. P values for pairwise comparisons are indicated in the graphs.

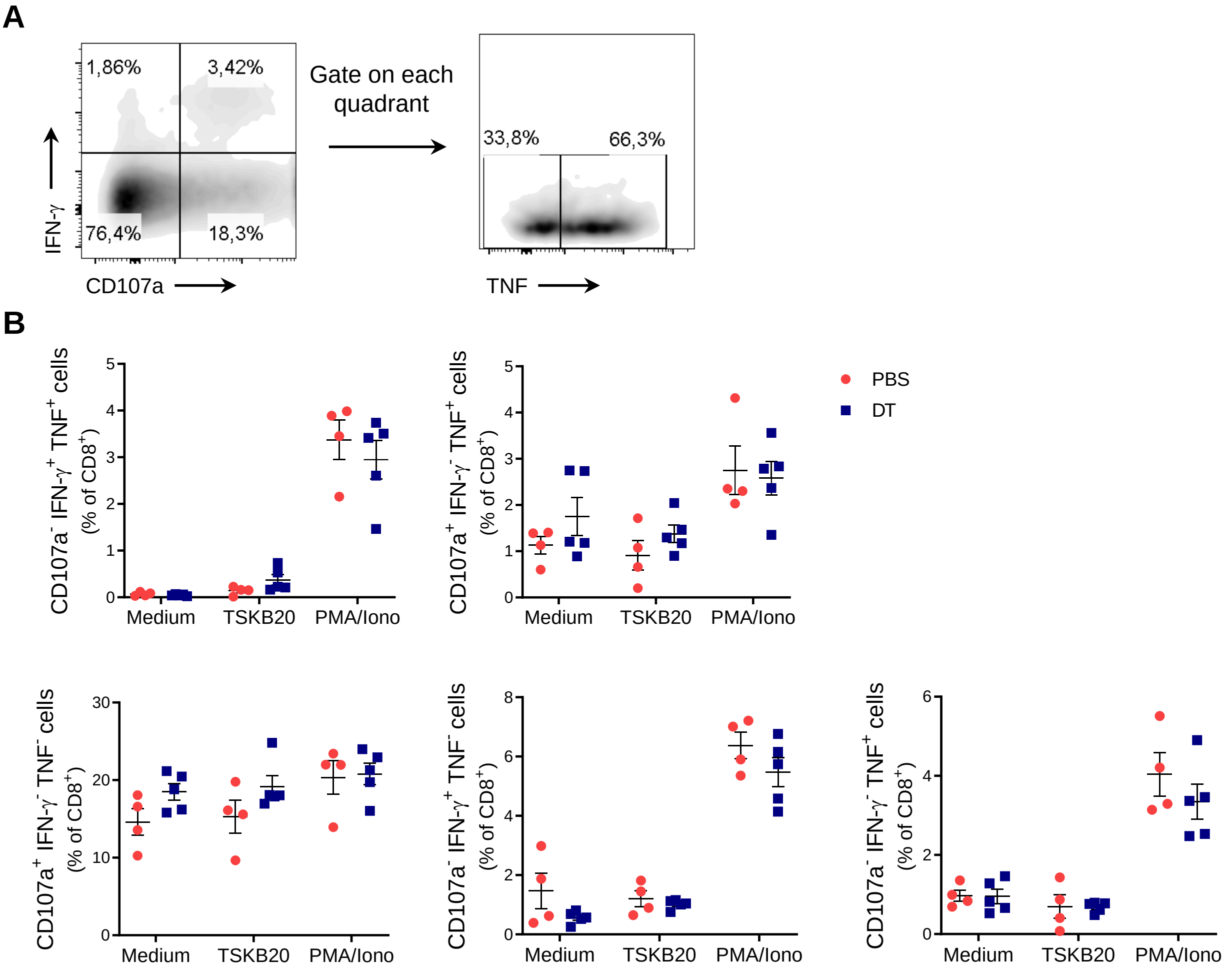


**Supplemental Figure 4. Effect of early Treg cell depletion on CD8+ T cell function. A)** Gating strategy for assessing effector cytokine production and CD107a surface mobilization in gated CD8+ cells. **B)** Percentage of CD8+ T cells from the spleen of PBS or DT-treated DEREG mice at day 21 pi that exhibit different combinations of effector functions, including CD107a mobilization and/or IFN-γ and/or TNF production upon 5h of the indicated stimulation. Medium condition was used as a negative control, while PMA/Ionomycin (PMA/Iono) was used as a positive control for polyclonal CD8+ T cell stimulation. Similar results were obtained in 2 independent experiments. All data are presented as mean ± SEM. Each symbol represents one individual mouse. Statistical significance was determined by Mann Whitney test. ns = not significant.

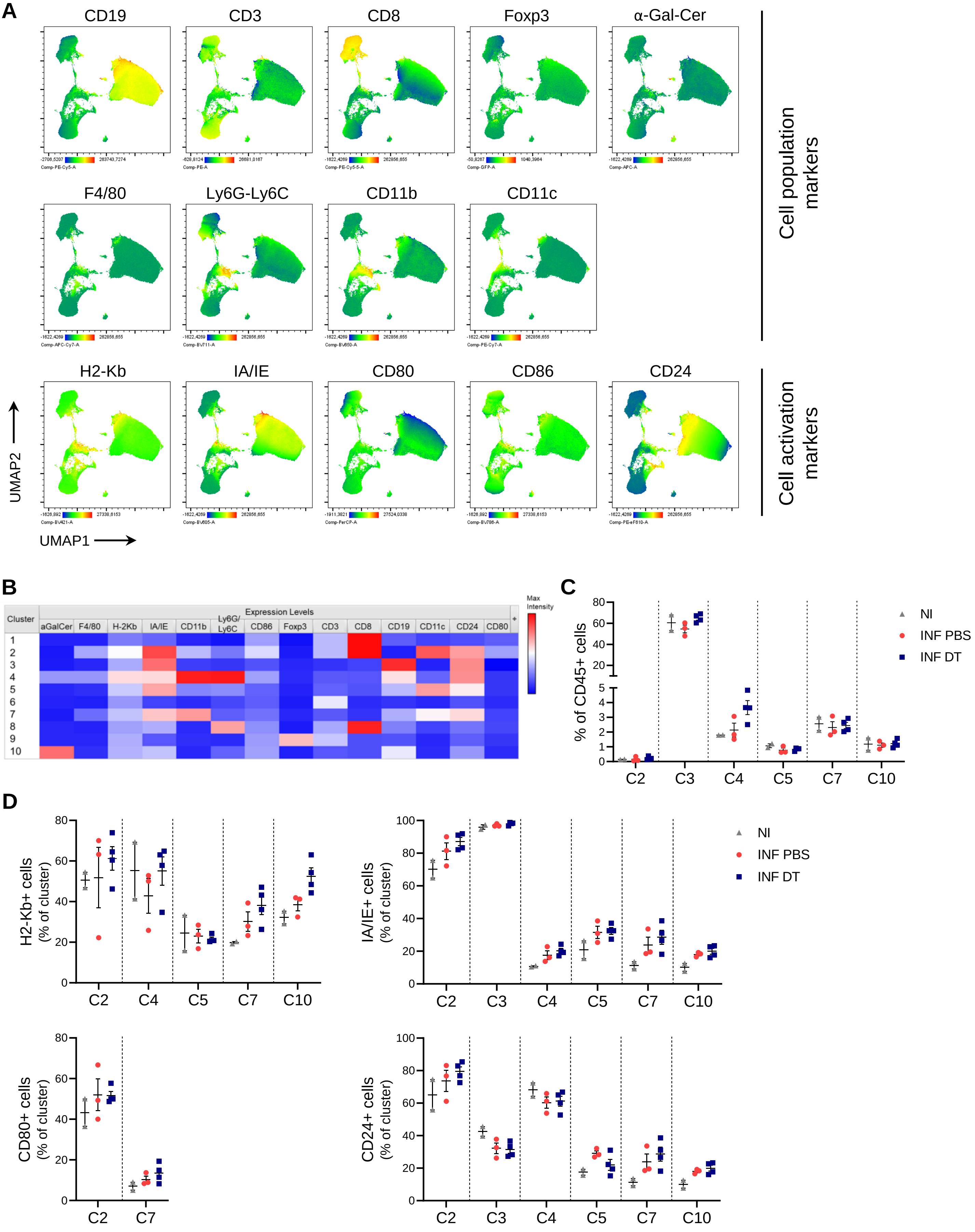
A
CD19
CD3
CD8
Foxp3
α-Gal-Cer
F4/80
Ly6G-Ly6C
CD11b
CD11c
H2-Kb
IA/IE
CD80
CD86
CD24
Cell population markers
Cell activation markers
UMAP2
UMAP1
B
Cluster
Expression Levels
aGalCer
F4/80
H-2Kb
IA/IE
CD11b
Ly6G/ Ly6C
CD86
Foxp3
CD3
CD8
CD19
CD11c
CD24
CD80
Max Intensity
C
% of CD45+ cells
NI
INF PBS
INF DT
C2
C3
C4
C5
C7
C10
D
H2-Kb+ cells (% of cluster)
IA/IE+ cells (% of cluster)
CD80+ cells (% of cluster)
CD24+ cells (% of cluster)

**Supplemental Figure 5. Frequencies and expression of activation markers in APC and innate cell clusters. A)** UMAP visualization for splenocytes expression of the different population and activation markers used in the flow cytometry panel for APC and innate cells characterization. Samples from the three experimental groups (NI, INF PBS and INF DT) are shown together. **B)** Heat map showing the expression level of each marker in the different clusters. **C)** Frequencies of selected clusters in total leukocytes (CD45+ cells) from the spleen of PBS or DT-treated DEREG mice at day 7 pi and non-infected controls. **D)** Frequency of cells expressing the indicated markers in the different clusters defined in Fig 4A. Clusters without positive cells for the corresponding marker were excluded from the analysis. All data are presented as mean ± SEM. Each symbol represents one individual mouse. Similar results were obtained in 3 independent experiments.

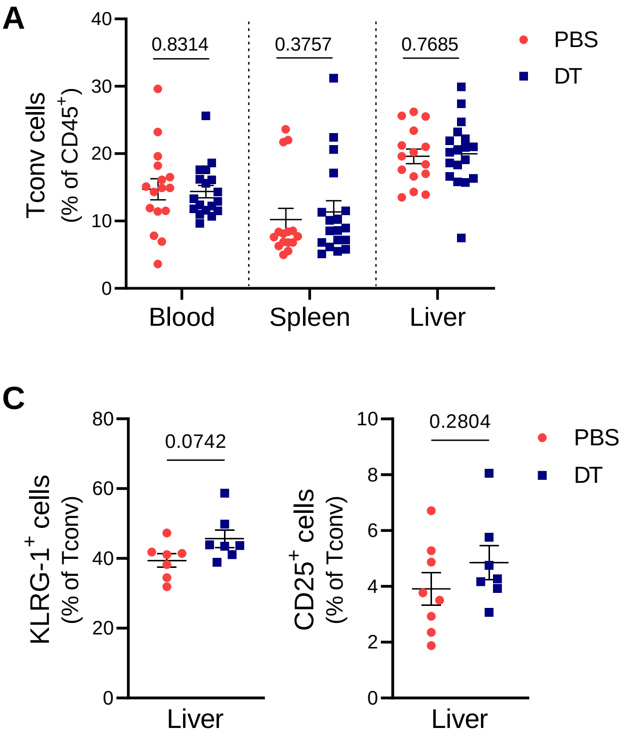


**Supplemental Figure 6. Effect of early Treg cell depletion on Tconv cell response. A-B)** Frequencies (A) and absolute numbers (B) of Tconv cells in blood, spleen and liver from PBS or DT-treated DEREG mice at day 20 pi. C) Frequency of KLRG-1+ and CD25+ Tconv cells in the liver of PBS or DT-treated DEREG mice at day 11 pi. All data are presented as mean ± SEM. Each symbol represents one individual mouse. Data were pooled from 2-4 independent experiments. Statistical significance was determined by Unpaired t test or Mann Whitney test, according to data distribution. P values for pairwise comparisons are indicated in the graphs.

**Supplementary Table 1. List of antibodies used for flow cytometry.**

| Target | Clone | Conjugate | Brand |
|---|---|---|---|
| CD107a | 1D4B | PE | Biolegend |
| CD11b | M1/70 | Super Bright 645 | eBioscience |
| CD11c | N418 | PE-Cyanine7 | eBioscience |
| CD127 | eBioSB/199 | PerCP-eFluor 710 | eBioscience |
| CD127 | A7R34 | PE | eBioscience |
| CD19 | eBio1D3 | APC-eFluor 780, PE-Cyanine5 | eBioscience |
| CD24 | M1/69 | PE-eFluor 610 | eBioscience |
| CD25 | PC61.5 | PE-Cyanine7, PE-eFluor 610 | eBioscience |
| CD39 | 24DMS1 | Alexa Fluor 700, PE-Cyanine7, PerCP-eFluor 710, eFluor 660 | eBioscience |
| CD39 | Duha59 | PE-Dazzle 594 | Biolegend |
| CD3e | 145-2C11 | PE | eBioscience |
| CD4 | GK1.5 | PE, APC-eFluor 780, PerCP-eFluor 710, Super Bright 645, PE-Cyanine7 | eBioscience |
| CD4 | GK1.5 | Alexa Fluor 700, APC | Biolegend |
| CD40 | 1C10 | PE-Cyanine5 | eBioscience |
| CD44 | IM7 | APC-eFluor 780, PE-Cyanine5 | eBioscience |
| CD44 | IM7 | PerCP-Cyanine5.5 | Biolegend |
| CD45 | 30-F11 | APC-Cyanine7 | BD Pharmingen |
| CD45 | 30-F11 | Alexa Fluor 700, PE-Cyanine7 | eBioscience |
| CD62L | MEL-14 | PerCP-Cyanine5.5, Super Bright 600 | eBioscience |
| CD69 | H1-2F3 | PE | BD Biosciences |
| CD69 | H1.2F3 | APC-Cyanine7 | Biolegend |
| CD80 | 16-10A1 | PerCP-eFluor 710 | eBioscience |
| CD86 | GL-1 | Brilliant Violet 785 | Biolegend |
| CD8a | 53-6.7 | PE, PerCP-Cyanine5.5, PE-Cyanine7, Alexa Fluor 700, PE-Cyanine5.5 | eBioscience |
| CTLA-4 | UC10-4B9 | Brillant Violet 605 | Biolegend |
| CTLA-4 | UC10-4B9 | APC | eBioscience |
| F4/80 | BM8 | APC-Cyanine7 | Biolegend |
| Foxp3 | FJK-16s | PerCP-Cyanine5.5, FITC | eBioscience |
| GITR | DTA-1 | Super Bright 600 | eBioscience |
| IFN-$\gamma$ | XMG1.2 | Brilliant Violet 711 | Biolegend |
| Ki-67 | SolA15 | eFluor 660 | eBioscience |
| KLRG-1 | 2F1 | PE | Biolegend |
| KLRG-1 | 2F1 | PE-eFluor 610 | eBioscience |
| LAG-3 | C9B7W | PerCP-Cyanine5.5 | Biolegend |
| LAG-3 | C9B7W | Brilliant Violet 421 | BD Biosciences |
| Ly-6C | HK1.4 | APC | eBioscience |
| Ly-6G/Ly-6C | RB6-8C5 | Super Bright 702 | eBioscience |
| MHC Class I (H-2kb) | AF6-88.5.5.3 | Super Bright 436 | eBioscience |
| MHC Class II (I-A/I-E) | M5/114.15.2 | Super Bright 600 | eBioscience |
| PD-1 | J43 | PE-Cyanine7 | eBioscience |
| PD-1 | 29F.1A12 | Brilliant Violet 421 | Biolegend |
| TIM3 | RMT3-23 | PE | eBioscience |
| TNF | MP6-XT22 | PerCP-Cyanine5.5 | Biolegend |
| TNF | MP6-XT22 | PE | eBioscience |
